# Cytokinin signaling in *Mycobacterium tuberculosis*

**DOI:** 10.1101/168815

**Authors:** Marie I. Samanovic, Hao-Chi Hsu, Marcus B. Jones, Victoria Jones, Michael R. McNeil, Samuel H. Becker, Ashley T. Jordan, Miroslav Strnad, Changcheng Xu, Mary Jackson, Huilin Li, K. Heran Darwin

## Abstract

It was reported the human-exclusive pathogen *Mycobacterium (M.) tuberculosis* secretes cytokinins, which previously had only been known as plant hormones. Cytokinins are adenine-based signaling molecules in plants that have never been shown to participate in signal transduction in other kingdoms of life. Here, we show that cytokinins induce the strong expression of the *M. tuberculosis* gene, Rv0077c. We found that a TetR-like transcriptional regulator, Rv0078, directly repressed expression of the Rv0077c gene. Strikingly, cytokinin-induced expression of Rv0077c resulted in a loss of acid-fast staining of *M. tuberculosis*. Although acid-fast staining is thought to be associated with changes in the bacterial cell envelope and virulence, Rv0077c-induced loss of acid-fastness did not affect antibiotic susceptibility or attenuate bacterial growth in mice. Collectively, these findings show cytokinins signal transcriptional changes that affect the *M. tuberculosis* cell envelope, and that cytokinin signaling is no longer limited to the kingdom plantae.

## Introduction

*M. tuberculosis* is the causative agent of tuberculosis, one of the world’s leading causes of mortality(WHO, 2017). For this reason, researchers are eager to identify pathways that could be targeted for the development of new therapeutics to treat this devastating disease. Among the current prioritized targets is the mycobacterial proteasome. *M. tuberculosis* strains with defects in proteasome-dependent degradation are highly attenuated in mice, partly because they are sensitive to nitric oxide (NO) (Cerda-Maira *et al.*, 2010, Darwin *et al.*, 2003, Gandotra *et al.*, 2010, Gandotra *et al.*, 2007, Lamichhane *et al.*, 2006, Lin *et al.*, 2009). The NO-sensitive phenotype of mutants defective for proteasomal degradation has been attributed to a failure to degrade an enzyme called Log (<u>Lo</u>nely <u>g</u>uy), a homologue of a plant enzyme involved in the synthesis of a family of *N^6^*-substituted adenine-based molecules called cytokinins (Samanovic *et al.*, 2015). The accumulation of Log in *M. tuberculosis* results in a buildup of cytokinins, a break down product of which includes aldehydes that effectively sensitize mycobacteria to NO (Samanovic *et al.*, 2015).

While we determined that a lack of proteasome-dependent degradation results in cytokinin accumulation, we were left with more questions, namely, what is the function of cytokinin production by *M. tuberculosis?* In plants, cytokinins are hormones that regulate growth and development (Sakakibara, 2006). In addition, bacterial plant pathogens and symbionts use cytokinins to facilitate the parasitism of their plant hosts (Frebort *et al.*, 2011). Outside of the laboratory, *M. tuberculosis* exclusively infects humans and is not known to have an environmental reservoir; therefore, it is unlikely *M. tuberculosis* secretes cytokinins to modulate plant development. Instead, we hypothesized that *M. tuberculosis*, like plants, uses cytokinins to signal intra-species transcriptional changes to its benefit. Here, we show that cytokinins induce the transcription of a gene of unknown function. Moreover, we identified and characterized a TetR-like regulator that represses the expression of this gene. While we have not yet identified an *in vivo* phenotype associated with this cytokinin-inducible gene, we found that its expression altered the cell envelope of *M. tuberculosis*, changing its staining properties. Collectively, these studies provide a foundation to characterize cytokinin signaling in *M. tuberculosis* and other cytokinin-producing bacterial species.

## Results

### Cytokinins induce the specific and high expression of Rv0077c in *M. tuberculosis*

To test if a cytokinin (CK) could induce gene expression in *M. tuberculosis*, we grew wild type (WT) *M. tuberculosis* H37Rv to mid-logarithmic phase and incubated the bacteria for five hours with *N^6^*-(Δ^2^-isopentenyl)adenine (iP), one of the most abundantly produced cytokinins in *M. tuberculosis* that is also commercially available(Samanovic *et al.*, 2015) (**see Materials and Methods--table supplement 1**). Using RNA sequencing (RNA-Seq), we discovered the expression of four genes, Rv0076c, Rv0077c, Rv0078, and *mmpL6*, was significantly induced upon iP treatment compared to treatment with the vehicle control (dimethylsulfoxide, DMSO) (**Figure 1A--table supplement 2**). Rv0077c is conserved among many mycobacterial species, while Rv0076c and Rv0078 are present only in several mycobacterial genomes (**Figure 1B**) (Lechat *et al.*, 2008). Notably, *M. smegmatis*, a distant, non-pathogenic relative of *M. tuberculosis*, has a weak homologue of Rv0077c and no conspicuous Rv0078 homologue (**Figure 1B**). *mmpL6* is one of 13 *mmpL* (mycobacterial membrane protein large) genes in *M. tuberculosis*. In strain H37Rv, *mmpL6* is predicted to encode a 42 kD protein with five trans-membrane-domains and is truncated compared to the same gene in ancestral tuberculosis strains (Brosch *et al.*, 2002). Thus, it is unclear if *mmpL6* encodes a functional protein in strain H37Rv.

**Fig. 1.**
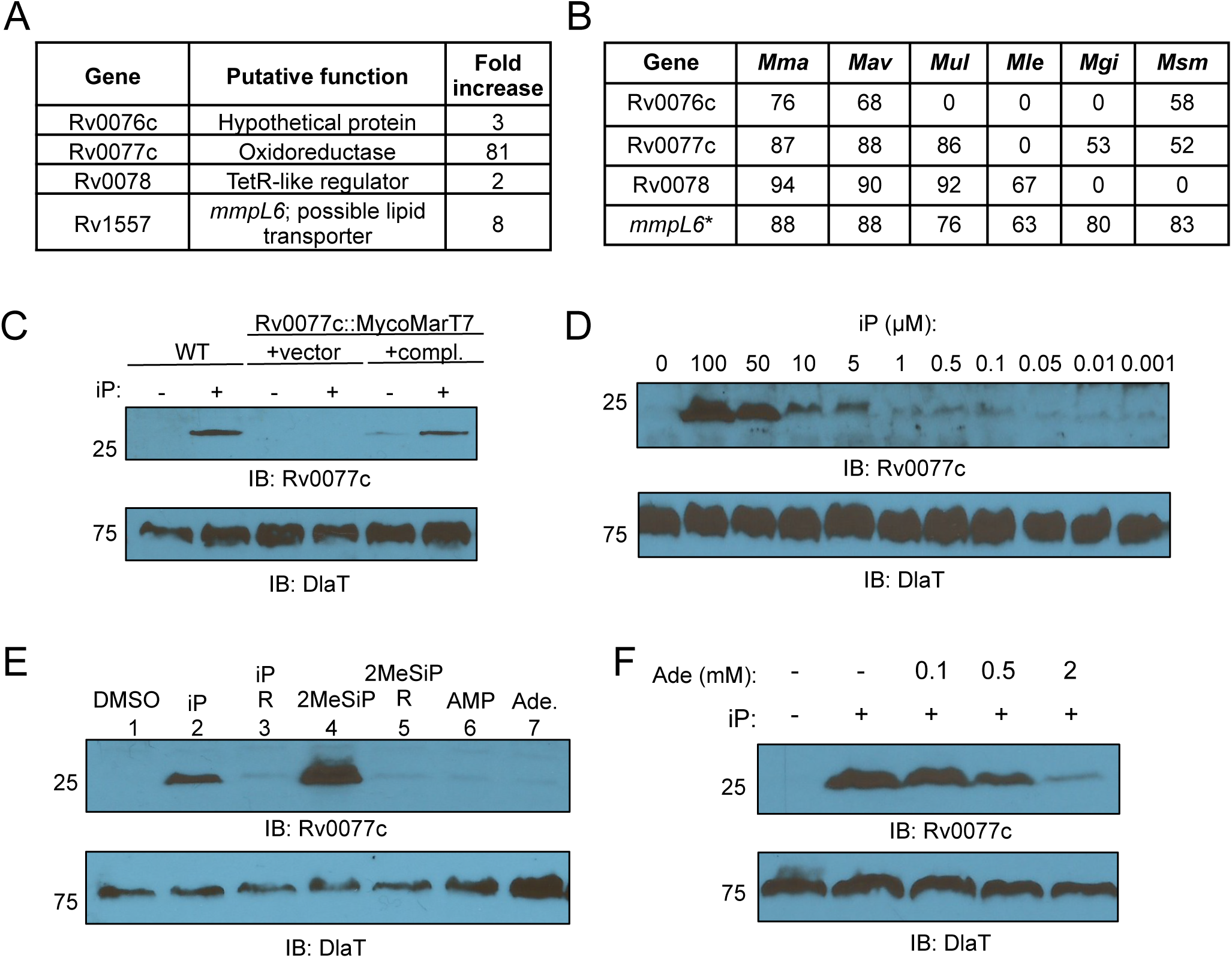
Cytokinins induce the expression of Rv0077c in M. tuberculosis. **(A)** Genes significantly regulated by the presence of 100 μM of iP for five hours as analyzed by RNA-Seq. **(B)** Percent identity between M. tuberculosis H37Rv proteins and proteins of targeted mycobacterial genomes including M. marinum (Mma), M. avium (Mav), M. ulcerans (Mul), M. gilvum (Mgi), and M. smegmatis (Msm). Asterisk (*) indicates mmpL6 encodes a truncated protein in H37Rv, unlike in the other mycobacterial species in the table. **(C)** Immunoblot for Rv0077c in total cell lysates of WT M. tuberculosis, “compl.” = complemented. iP was used at a final concentration of 100 μM when added. **(D)** Dose-dependent production of Rv0077c protein. Bacteria were incubated with cytokinin at the indicated concentrations for 24 hours. **(E)** Only cytokinins, and not closely related molecules, induce the production of Rv0077c. Each compound was added to a final concentration of 100 μM. “R” indicates the riboside form of the preceding indicated cytokinin. **(F)** Adenine inhibits the induction of Rv0077c by 100 μM iP. For all panels, we added an equal volume of DMSO to samples where iP was not added. For all immunoblots (IB), we stripped the membranes and incubated them with antibodies to dihydrolipoamide acyltransferase (DlaT) to confirm equal loading of total lysates. Molecular weight standards are indicated to the left of the blots and are in kilodaltons (kD). Ade, adenine.

Rv0077c was by far the most strongly induced gene in *M. tuberculosis* upon iP treatment therefore we chose it for follow-up studies. We raised polyclonal antibodies to recombinant Rv0077c protein and showed that protein levels were increased in *M. tuberculosis* treated with iP for 24 hours (**Figure 1C,** first two lanes). Rv0077c was barely detectable in cell lysates of bacteria that had not been incubated with iP and was undetectable in a strain with a transposon insertion mutation in Rv0077c (**Figure 1C,** center two lanes). Rv0077c protein was restored to WT levels in the mutant upon complementation with an integrative plasmid encoding Rv0077c expressed from its native promoter (**Figure 1C,** last two lanes). We also found a dose-dependent induction of Rv0077c production using iP concentrations from 1 nM to 100 µM (**Figure 1D**).

We next synthesized and tested if the most abundantly produced cytokinin in *M. tuberculosis*, 2-methyl-thio-iP (2MeSiP) (Samanovic *et al.*, 2015), could also induce Rv0077c production. 2MeSiP strongly induced Rv0077c production (**Figure 1E,** lane 4). Importantly, we did not observe induction of Rv0077c when we incubated the bacteria with the appropriate cytokinin riboside (R) precursors iPR or 2MeSiPR (**Figure 1E,** lanes 3 and 5). Similarly, adenosine monophosphate (AMP) or the closely related molecule adenine could not induce Rv0077c expression (**Figure 1E,** lanes 6 and 7). We hypothesized while adenine could not induce Rv0077c expression, at high enough concentrations it could possibly inhibit Rv0077c induction by competing with cytokinin for access to a transporter or receptor. Indeed, adenine reduced the induction of Rv0077c by iP in a dose-dependent manner (**Figure 1F**).

### Identification of an operator for the TetR-like transcriptional repressor Rv0078

Rv0077c is divergently expressed from Rv0078; the proposed translational start codons for these genes are separated by 61 base pairs therefore we hypothesized that Rv0078 encodes a repressor of Rv0077c expression. We identified the promoter regions for each gene by performing rapid amplification of 5′ complementary DNA ends (5′RACE) analysis for Rv0077c and Rv0078 and determined the likely start of transcription for one gene was within the 5′ untranslated region of the other gene (**Figure 2A**). We sought to identify an operator sequence by using an electrophoretic mobility shift assay (EMSA, **Figure 2B**). We narrowed down a putative Rv0078 binding site to a region overlapping the proposed starts of transcription (+1) of both Rv0077c and Rv0078 (**Figure 2A**, red box). TetR-like regulators generally bind to inverted repeat sequences; we identified a 21-base pair (bp) sequence within −14/+19 of Rv0077c containing two 10-bp inverted repeat sequences. Mutagenesis of the probe in the repeat sequences disrupted Rv0078 binding to the DNA (**Figure 2B**). Importantly, the binding sequence overlaps the putative transcriptional start sites of both genes, suggesting that expression of both genes is repressed by Rv0078. Notably, the addition of iP did not result in the release of Rv0078 from the DNA probe (**Figure 2B--figure supplement 1**). Therefore, cytokinins do not appear to directly bind to Rv0078 to induce gene expression.

**Fig. 2.**
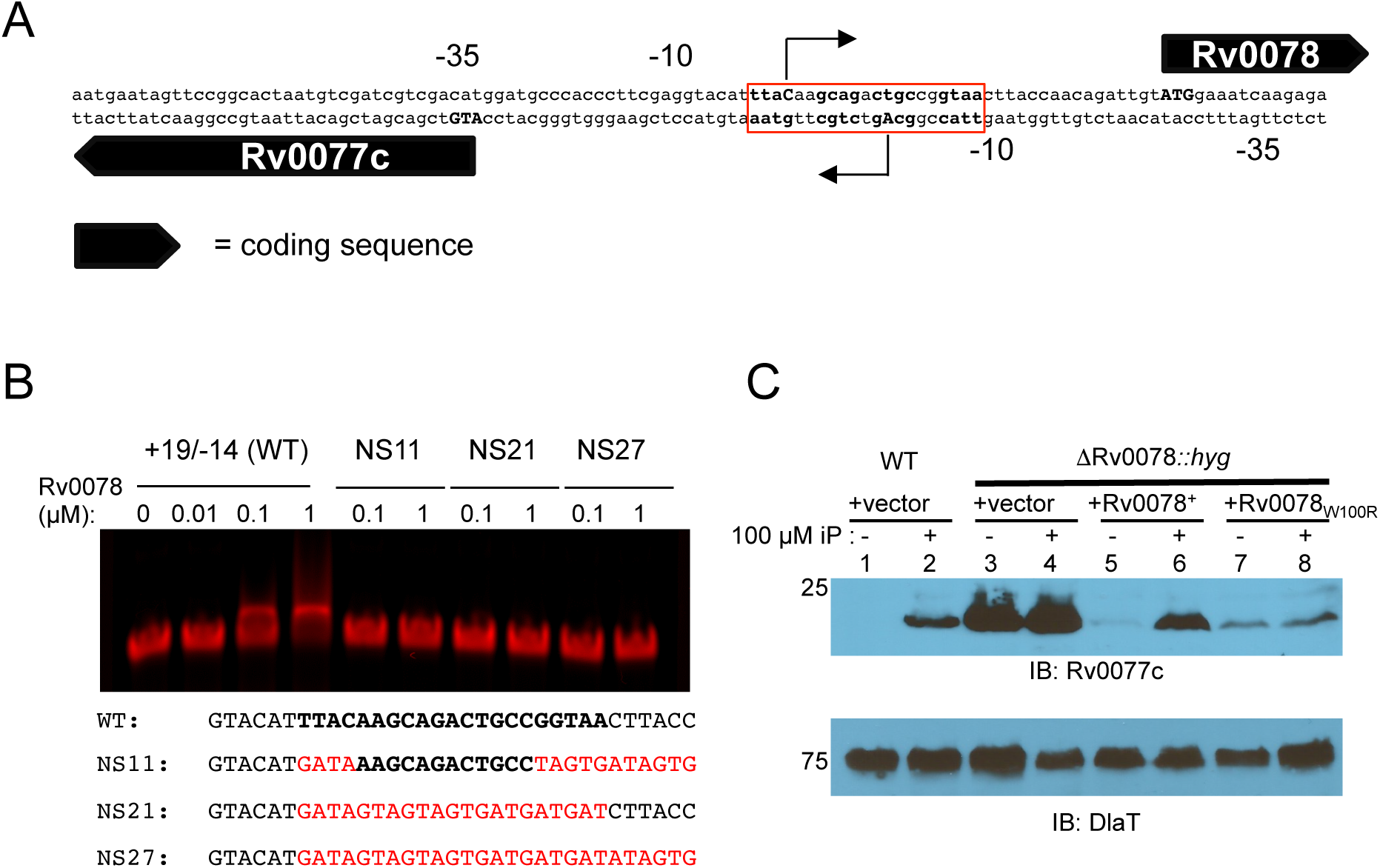
Rv0078 represses the expression of Rv0077c. **(A)** The putative transcriptional start sites (+1) of Rv0077c and Rv0078 as determined by 5′RACE and represented as bent arrows. The predicted start codons are in capital bold letters. **(B)** EMSA analysis identifies a putative repressor binding site. Probe sequences. +19/-14 refers to positions relative to the Rv0077c +1. In bold is the presumed binding site. Mutated residues are in red. Not shown at the end of each probe is a sequence for annealing to a fluorescent tag (Table supplement 1). Rv0078 was purified under native conditions from *E. coli.* **(C)** Deletion and disruption of Rv0078 results in the constitutive expression of Rv0077c. Total cell lysates were prepared and separated on a 10% SDS-PAGE gel. IB = immunoblot. The membrane was stripped and incubated them with antibodies to DlaT to confirm equal loading of samples. Molecular weight standards are indicated to the left of the blots and are in kilodaltons (kD).

Based on the 5′RACE analysis we were able to delete and replace most of the Rv0078 gene with the hygromycin-resistance gene (*hyg*) without disrupting the promoter of Rv0077c in *M. tuberculosis* H37Rv. A ΔRv0078∷*hyg* strain displayed constitutively high expression of Rv0077c irrespective of the presence of cytokinin, supporting a model where Rv0078 directly represses Rv0077c expression (**Figure 2C**). A single copy of Rv0078 expressed from its native promoter restored iP-regulated control of Rv0077c in this strain (**Figure 2C,** lanes 5 and 6). Interestingly, in the process of making the complementation plasmid, we acquired a random mutation (likely generated during PCR) in Rv0078 that changed a tryptophan to arginine (W100R); this allele was unable to complement the ΔRv0078∷*hyg* strain (**Figure 2C,** lanes 7 and 8).

### Two Rv0078 dimers bind to one operator

To gain an understanding of how Rv0078 represses gene expression, we solved the crystal structure of Rv0078 to 1.85 Å by single-wavelength anomalous dispersion method. As previously reported, Rv0078 forms dimers resembling other TetR-like proteins (Orth *et al.*, 2000, Wohlkonig *et al.*, 2017). Unlike the canonical TetR binding site, *tetO*, which is 15 bp long, the Rv0078 binding site is 21 bp long suggesting Rv0078 binds to DNA differently than TetR. We performed an isothermal titration calorimetry (ITC) experiment and determined that the apparent *K_D_* of Rv0078 for operator DNA was 357 nM (**Figure 3A**). The estimated stoichiometry of DNA duplex to Rv0078 dimer was 0.66, suggesting that two Rv0078 dimers bind to one operator sequence.

**Fig. 3.**
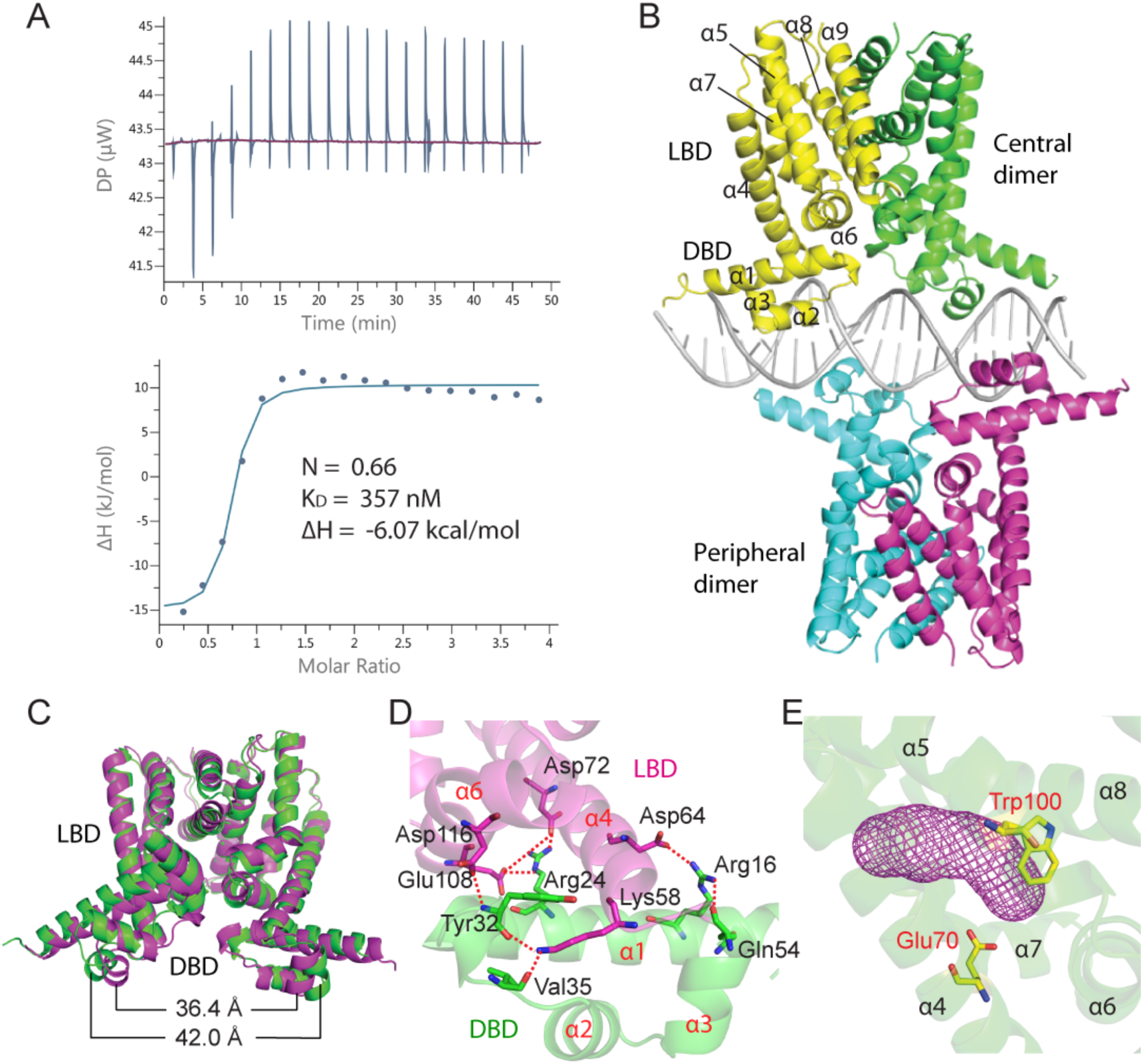
Crystal structure of Rv0078 in complex with DNA. **(A)** ITC of Rv0078 binding to the +13/-8 DNA probe. The binding stoichiometry, ΔH, and K_D_ are marked. **(B)** Overall structure of Rv0078-DNA complex in cartoon view. Two Rv0078 dimers (“central” and “peripheral”) bind to one DNA molecule. **(C)** The distance between two DNA-binding domains decreases by ~6 Å when bound to DNA. The DNA-free Rv0078 is in green; the DNA-bound Rv0078 is in magenta. **(D)** Interactions between the LBD (magenta) and DBD (green) in a monomer. **(E)** The ligand binding pocket (magenta mesh) of Rv0078 is enclosed by a four-helix bundle (helices α5 to α8).

We next co-crystallized Rv0078 with a 23-bp DNA fragment that was extended one base pair at each end of the 21-bp operator sequence identified in Figure 2 (−8/+13 of Rv0077c) and solved the structure to 3.0 Å resolution (**Table supplement 3**). As predicted by the ITC experiments, we observed two Rv0078 dimers bound to a DNA duplex where each dimer bound on opposite sides of the DNA (**Figure 3B**). The top or “central” dimer bound the DNA palindrome symmetrically while the bottom or “peripheral” dimer bound to DNA off-center, staggered from the central dimer binding site by seven base pairs. Thus, the longer-than-canonical (i.e., *tetO*) binding site is required for accommodating two Rv0078 dimers, unlike TetR, which binds to *tetO* as a single dimer.

Rv0078 has an N-terminal DNA-binding domain (DBD) and a C-terminal ligand-binding domain (LBD) (**Figure 3B-D**). The DBD is comprised of three α-helices (α1 - α3), and within this domain, α2 and α3 form a helix-turn-helix (HTH) motif that directly contacts DNA. The LBD is formed by six α-helices (α4 - α9). The four Rv0078 protein structures within a complex were nearly the same; the root mean square deviation (rmsd) ranged from 0.07 Å to 0.17 Å in a pair-wise superimposition. The Rv0078 dimer structure is held together by α8 and α9 of each monomer, forming a four-helix bundle. The two dimer structures were highly similar with an rmsd of 0.14 Å. DNA binding induced significant conformational changes across the Rv0078 dimer structure, with an rmsd of 2.13 Å when compared to the DNA-free dimer. In particular, the two α3 helices move towards each other by ~6 Å, in order to reduce their distance to 36.4 Å and fit in the DNA major grooves (**Figure 3C**). In the LBD, the ligand entry between helices α4 and α5 is open and the ligand-binding pocket is empty. It appears that changes initiated in the DBD regions upon binding to the DNA are transmitted to the LBD via the DBD-LBD interface that involves extensive interactions, including two salt bridges and six H-bonds (**Figure 3D**).

The ligand-binding pocket of Rv0078 is largely hydrophobic. Within this pocket, we observed an elongated density resembling a long aliphatic chain of a fatty acid (**Figure supplement 2**). Gas chromatograph-mass spectrometry of the compounds extracted from Rv0078 purified from *E. coli* revealed fatty acids commonly found in this organism, and a palmitate molecule fit well with the electron density (**Figure supplement 2A-E**). A fatty acid carboxylate formed a hydrogen bond with Rv0078 Glu-70 (**Figure 3E--figure supplement 2C**), and the long alkyl chain had numerous hydrophobic interactions within the extended ligand pocket. When we purified Rv0078 under denaturing conditions and refolded the protein to remove the lipid, we observed the same EMSA results as we observed with protein purified under native conditions (data not shown), suggesting this fatty acid is unlikely to be the native Rv0078 ligand; however, these data may suggest that the native ligand is fatty acid-like. Interestingly, we found that Trp100 faces the ligand-binding pocket (**Figure 3E**), which suggests Trp100 interacts with the natural ligand. This hypothesis is supported by our data showing an Rv0078_W100R_ mutant was unable to complement the Rv0078-deletion mutant strain (**Figure 2C, lane 8**).

In addition to characterizing the LBD, we identified nine amino acids that were important for interacting with DNA (**Fig. 4A-E**). Specifically, the hydroxyl groups of Thr37, Thr47, and Tyr52 interacted strongly with DNA phosphates at a distance of 2.6 Å. Arg48 is the only residue that interacted with DNA by recognizing a guanine base at a distance of 2.6 Å. To further examine the importance of these residues, we introduced single amino acid substitutions (T37V, T47V, R48M, and Y52F) or double mutations (T47V, R48M) into Rv0078 and performed EMSA assays. While all of the mutant proteins were soluble and behaved like the WT protein in solution, Rv0078_R48M_ did not bind to DNA, and the other mutant proteins bound to DNA with reduced affinities (**Figure 4F**).

**Fig. 4.**
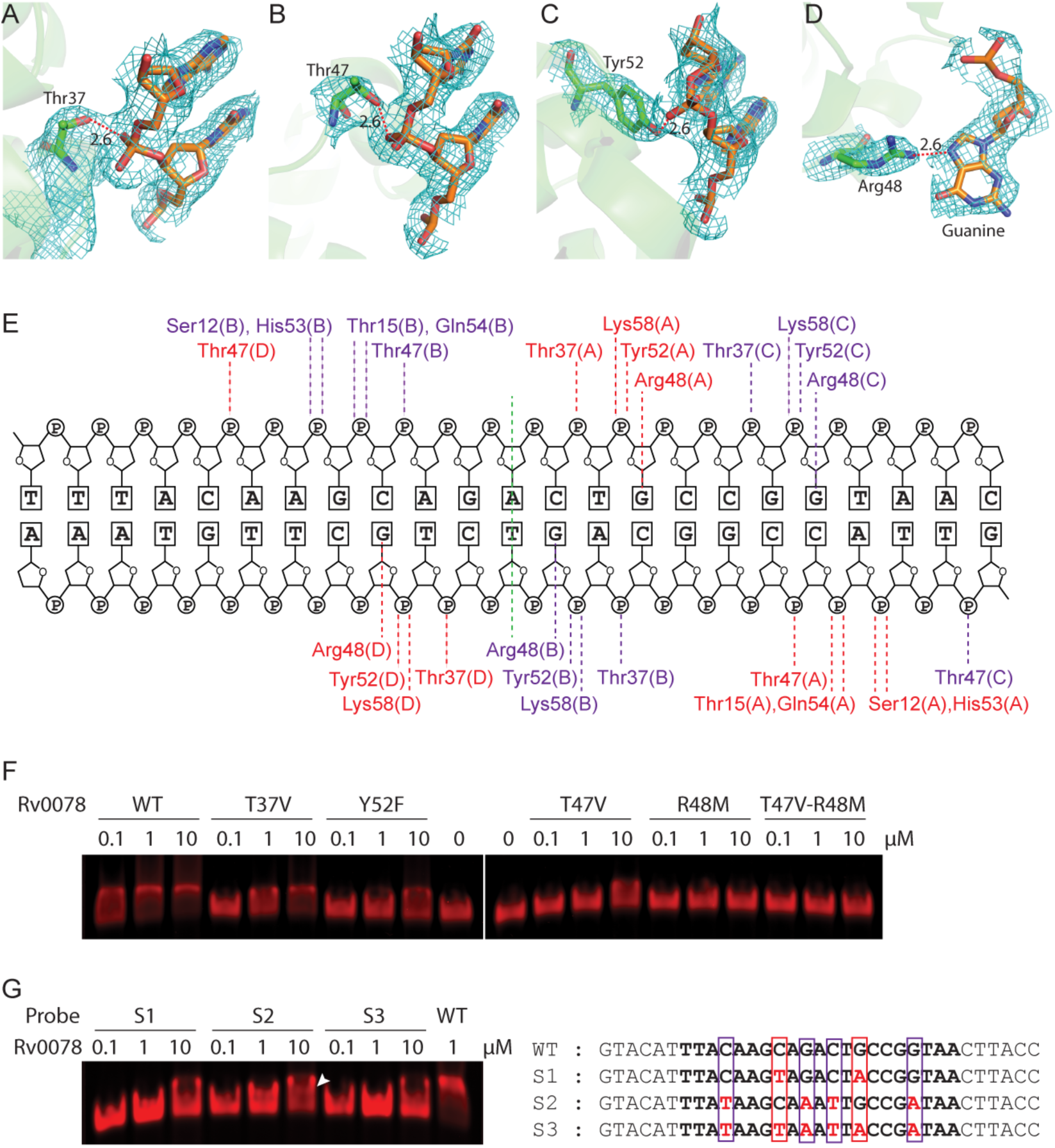
Rv0078-DNA interactions. The hydroxyl group of. **(A)** Thr37, **(B)** Thr47, and **(C)** Tyr52 interact with the backbone phosphate with a distance of 2.6 Å. **(D)** Arg48 interacts with guanidine with a distance of 2.6 Å. The 2Fo-Fc maps are contoured at 1σ level. **(E)** A schematic representation of Rv0078-DNA contacts. Residues of the central dimer are labeled in red, and residues of the peripheral dimer are in purple. **(F)** EMSA using WT DNA probe and Rv0078 with mutations in Thr37, Thr47, Arg48, or Tyr52. **(G)** Left panel: EMSA of Rv0078 with DNA probes with G-to-A substitutions in the central dimer binding region (S1), peripheral dimer binding region (S2), or in both DNA regions (S3). The white arrowhead marks the partial shift with the S2 probe at high protein concentration. Right panel: sequences of the four DNA probes used in EMSA experiments. The nucleotides contacting the central dimer are boxed in red and those contacting the peripheral dimer are boxed in purple.

To examine the DNA sequence specificity of Rv0078, we synthesized three EMSA probes by changing the Rv0078 Arg48-interacting guanines to adenines (G-to-A) in the central dimer binding region (probe S1), the peripheral dimer-binding region (probe S2), or both dimer-binding regions (probe S3). These G-to-A substitutions either abolished or compromised Rv0078 binding to DNA, affirming the critical role of guanines in the binding site (**Figure 4G**). Notably, substitutions in probe S1 entirely abolished Rv0078 binding, while substitutions in S2 retained partial gel shift at a high concentration (white arrowhead in **Figure 4G**). This observation suggests that the binding of the two Rv0078 dimers is cooperative, with the central dimer likely the first one to bind to DNA.

### Constitutive expression of Rv0077c does not affect antibiotic susceptibility or virulence in mice

During our ongoing studies, a report was published on the identification of a small molecule of the spiroisoxazoline family, SMARt-420, which strongly induces the expression of the Rv0077c orthologue *bcg_0108c* in *M. bovis* Bacille Calmette-Guerin (Blondiaux *et al.*, 2017). Using x-ray crystallography and surface plasmon resonance techniques, the authors of this study found SMARt-420 binds to Rv0078 to derepress binding from the Rv0077c promoter (Blondiaux *et al.*, 2017, Wohlkonig *et al.*, 2017). SMARt-420 was identified in a search for compounds that boost the efficacy of the second-line tuberculosis drug ethionamide (ETH). ETH is a pro-drug that is activated by the mono-oxygenase EthA, which transforms ETH into highly reactive intermediates. Activated ETH and nicotinamide adenine dinucleotide form a stable adduct, which binds to and inhibits InhA, an essential enzyme needed for mycolic acid synthesis in mycobacteria (DeBarber *et al.*, 2000, Vannelli *et al.*, 2002). While spontaneous inactivating mutations in *ethA* result in resistance to ETH, it was proposed that the induction of Rv0077c expression could bypass the need for EthA and transform ETH into its toxic form (Blondiaux *et al.*, 2017). Based on this study, we predicted that an Rv0078 mutant of *M. tuberculosis*, which expresses high levels of Rv0077c, should be hypersensitive to ETH compared to the parental strain H37Rv. However, we observed either little to no significant change in the 50% minimum inhibitory concentration (MIC_50_) of ETH between the WT and ΔRv0078∷*hyg* strains (**Table supplement 4**). We also tested if the constitutive expression of Rv0077c changed the susceptibility of *M. tuberculosis* to other antibiotics, including two cell wall synthesis inhibitors. We observed no differences in the MIC_50_ of these antibiotics between the WT and ΔRv0078::*hyg* strains (**Table supplement 4**).

We also tested if either the Rv0077c or Rv0078 mutant had growth defects *in vivo* compared to the WT H37Rv strain. We infected C57BL/6J mice by a low-dose aerosol route with the parental, mutant, and complemented mutant strains, as well as with the Rv0078 mutant transformed with the Rv0078_W100R_ allele. Interestingly, none of the strains revealed a difference in growth or survival compared to WT *M. tuberculosis* in mice as determined by the recovery of colony forming units (CFU) from the lungs and spleens (**Figure 5--figure supplement 3**).

**Fig. 5.**
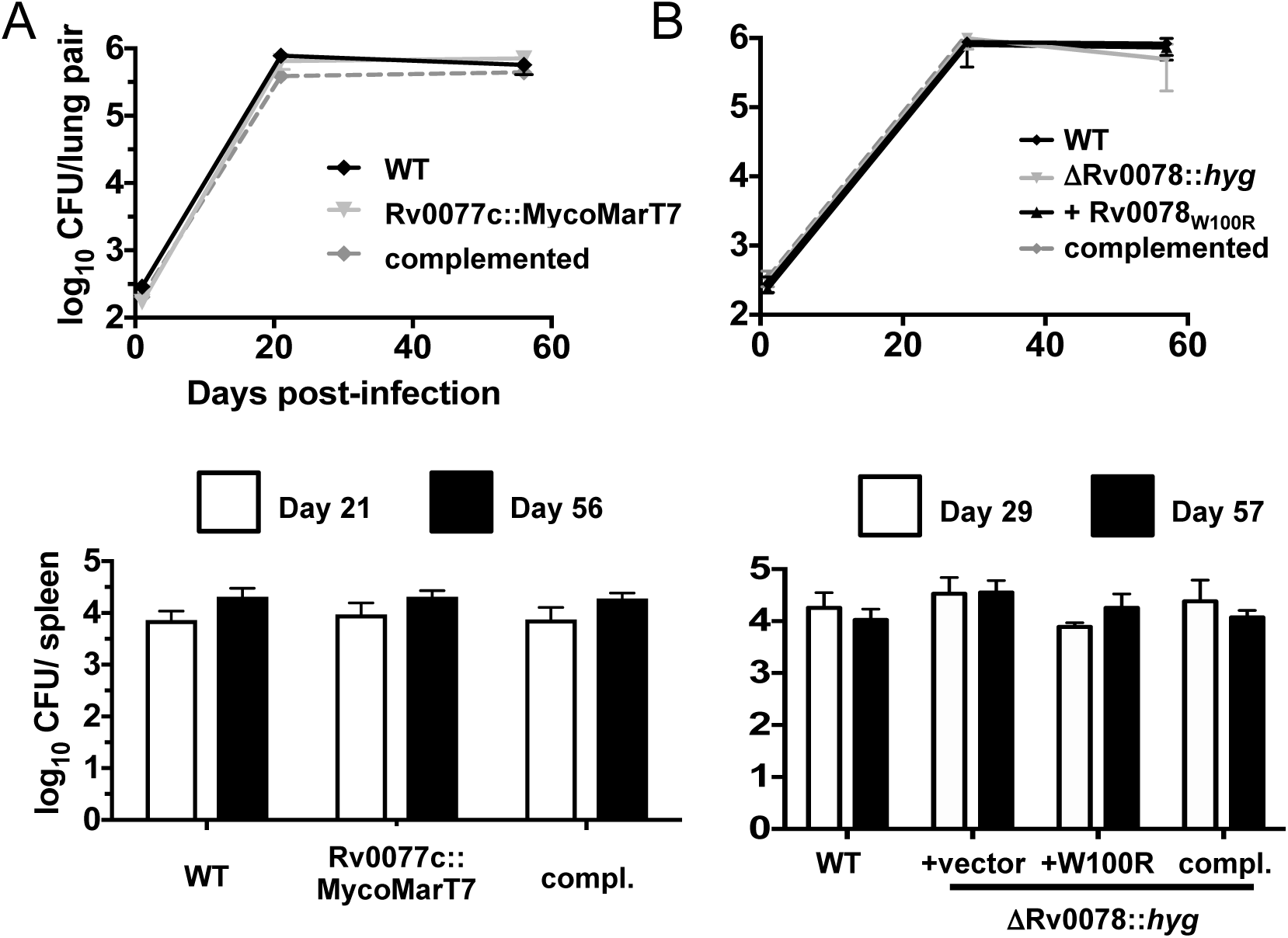
Loss of Rv0077c or Rv0078 does not attenuate bacterial survival in mice. **(A)** Bacterial colony forming units (CFU) after infection of C57BL/6J mice with WT, Rv0077c and complemented strains. **(B)** Bacterial CFU after infection of C57BL/6J mice with WT, Rv0078 and complemented strains. For both panels, data in each are from a single experiment that is representative of two independent experiments. Error bars indicate the standard error of the mean.

### Expression of Rv0077c alters acid-fast staining of M. tuberculosis

To gain insight into the function of Rv0077c, we performed metabolomics analysis of strains expressing Rv0077c to potentially determine how its presence alters bacterial physiology. We prepared total cell lysates of WT and Rv0077c mutant strains treated with or without 100 µM iP for 24 hours (see Materials and Methods). From a total of 337 detectable metabolites, we observed a significant change in 24 molecules after the addition of iP to WT *M. tuberculosis*. Seventeen metabolites showed a consistent difference between samples in which iP was added and these changes disappeared in an Rv0077c-disrupted strain, suggesting the changes were specifically due to the presence of Rv0077c (**Table supplement 4**, highlighted in yellow). We observed an increased abundance of several phospholipids and a decrease in a major precursor of peptidoglycan, N-acetyl-glucosamine-1-phosphate (GlcNAc1P). We therefore hypothesized that Rv0077c modified one or more components of the cell envelope. Microscopic examination of Ziehl-Neelsen (ZN) stained WT *M. tuberculosis* treated with iP showed a loss of acid-fast staining (**Figure 6**, panel a compared to b), and this phenotype depended on the presence of Rv0077c (**Figure 6**, panel c compared to d). Complementation of the mutation with Rv0077c alone restored the iP-induced loss of acid-fast staining (**Figure 6**, panels e and f). Deletion of Rv0078 resulted in a constitutive loss of staining, irrespective of the presence of iP (**Figure 6**, panels g and h). Complementation of the Rv0078 deletion with the WT gene restored iP-control of loss of acid-fast staining (**Figure 6**, panels i and j) but complementation with Rv0078_W100R_, which could not fully de-repress Rv0077c expression in the presence of iP (**Figure. 2C**), could not restore iP-induced loss of acid-fast staining (**Figure 6**, panels k and l). Mixing and simultaneous staining of the Rv0077c and Rv0078 mutants further showed the staining differences were not a result of a technical artifact (**Figure 6,** panel m).

**Fig. 6.**
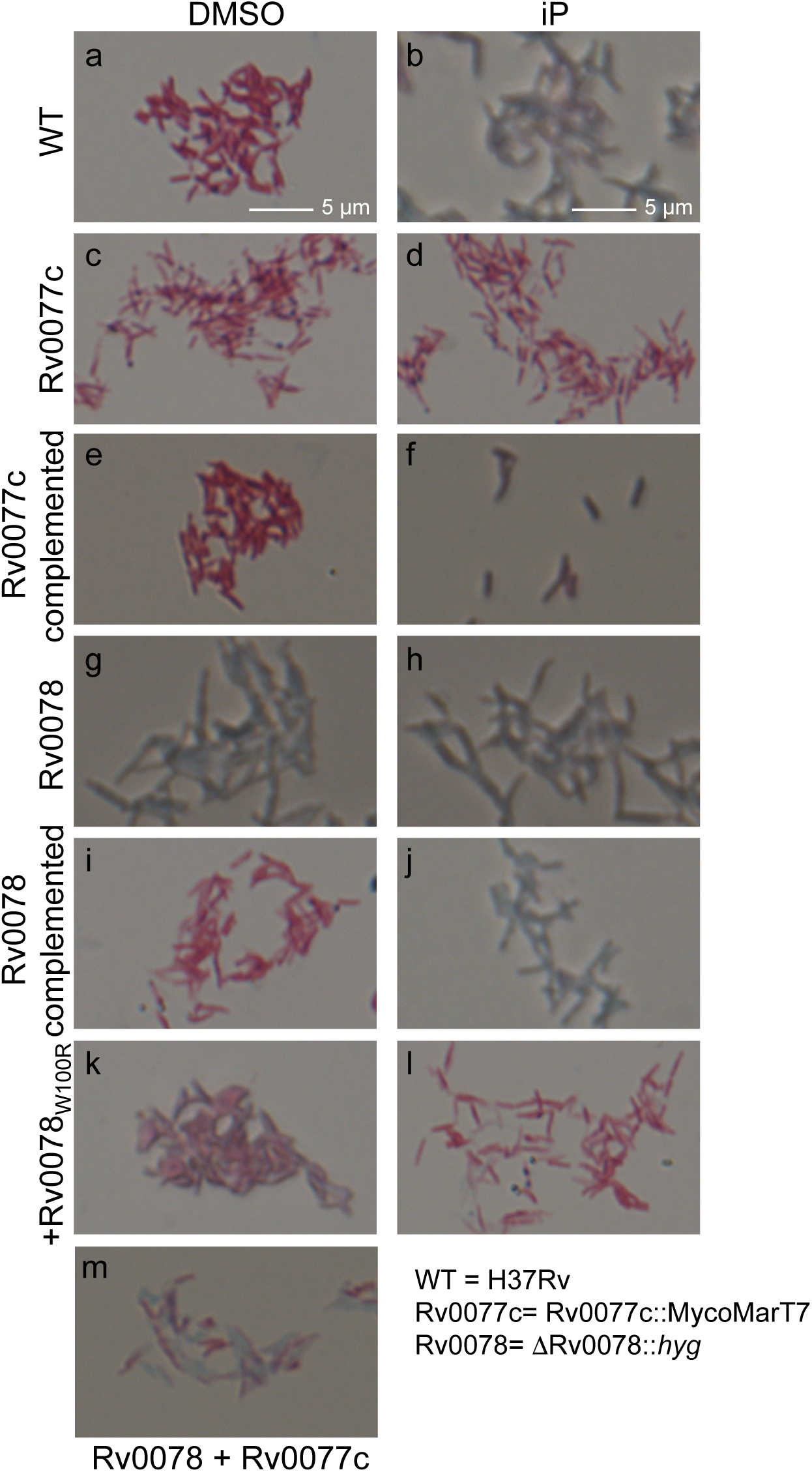
Induction of Rv0077c expression results in loss of acid-fast staining. *M. tuberculosis* strains were examined by staining. Magnification is 63-fold. Scale bars are applicable to all images. See text for details.

Acid-fast staining is primarily thought to be associated with mycolic acids; alterations in mycolic acid synthesis result in negative effects on cell growth *in vitro* and *in vivo*(Barkan *et al.*, 2009, Bhatt *et al.*, 2007). We used thin layer and liquid or gas chromatography coupled to mass spectrometry to analyze the fatty acid, mycolic acid and lipid contents of the WT, Rv0077c and Rv0078 strains and observed no significant qualitative or quantitative differences between strains (**Figure supplement 4, table supplement 6**). In particular, the expression of Rv0077c did not conspicuously modify the chain length of mycolic acids, their cyclopropanation or the relative abundance of keto- to alpha- mycolates, and did not alter the wax ester and triglyceride content of the strains that have been previously been linked to acid-fast staining (Bhatt *et al.*, 2007, Deb *et al.*, 2009).

## Discussion

Our studies are the first to demonstrate CKs induce robust and specific transcriptional and physiologic changes in a bacterial species. *M. tuberculosis* treated with CK expressed high levels of Rv0077c, which altered its metabolome and staining properties. Despite these changes, we did not observe an effect on the susceptibility of these bacteria to antibiotics of different classes or on survival in mice. In addition we found the transcriptional regulator Rv0078 represses Rv0077c expression in the absence of CK, and we defined the operator to which Rv0078 binds. We determined that two dimers of Rv0078 bind to inverted repeats in the intergenic region between Rv0077c and Rv0078, which likely represses the expression of both genes.

In plants, CKs are sensed by membrane receptors related to the sensors of bacterial two-component systems (TCS) (To & Kieber, 2008, Inoue *et al.*, 2001, Steklov *et al.*, 2013). In these systems, CKs interact with membrane receptors with a CHASE (<u>c</u>yclases/<u>h</u>istidine kinases <u>a</u>ssociated <u>s</u>ensing <u>e</u>xtracellular) domain (Mougel & Zhulin, 2001, Anantharaman & Aravind, 2001). As in bacteria, the ligand-receptor interaction stimulates a phospho-relay that ultimately leads to the phosphorylation of a response regulator protein, which either represses or activates gene expression. *M. tuberculosis* has at least 12 known TCS, and none has a predicted CHASE domain (L. Aravind, February 2012; and I. Jouline, August 2017, personal communications). Thus, it remains to be determined if a TCS sensor protein is involved in CK signal transduction in *M. tuberculosis*.

It also remains to be determined what the natural ligand is for Rv0078; while SMARt-420 binds robustly to Rv0078 (Wohlkonig *et al.*, 2017, Blondiaux *et al.*, 2017) the cytokinin iP did not in our studies. This result may not be surprising when considering SMARt-420 and cytokinins bear no resemblance to each other. Furthermore, we could not co-crystalize iP with Rv0078 (data not shown). We hypothesize that the interaction of CK with an unknown receptor or enzyme leads to the synthesis of a small molecule (e.g., a lipid) that binds to Rv0078, resulting in the derepression of Rv0077c. This possibility may be supported by our observation that there is an increase in several phospholipids in iP-treated *M. tuberculosis*. We are currently working to identify factors required for CK-mediated gene induction.

Rv0077c is predicted to have an α/β hydrolase fold (Soding *et al.*, 2005), which can have a variety of substrate types (Nardini & Dijkstra, 1999). Based on our metabolomics and microscopy studies, we predict Rv0077c targets one or more components of the cell envelope. The identification of the natural target of Rv0077c may help provide new insight into the elusive molecular basis of acid-fastness of this human-exclusive pathogen. Our results may also indicate that acid fast staining can be affected by different cell envelope chemistries in addition to changes in mycolic acid structure or content.

While we did not observe any differences in bacterial burden in mice infected with strains that either were disrupted for or constitutively expressed Rv0077c, it is possible that Rv0077c function might only be needed during very late or specific stages of infection. This idea may be supported by a previous observation that the cytokinin synthase *log* gene in *M. marinum* is specifically expressed in late granulomas of infected frogs (Ramakrishnan *et al.*, 2000). Furthermore, mice might not provide an optimal model to observe a role for this pathway in tuberculosis.

A previous report suggested the expression of Rv0077c increases the sensitivity of *M. tuberculosis* to the antibiotic ETH. ETH must be activated by a mono-oxygenase, EthA, in order for it to be toxic to *M. tuberculosis* (DeBarber *et al.*, 2000, Vannelli *et al.*, 2002). We found the expression of Rv0077c did not confer increased susceptibility to ETH or any other antibiotic we tested, an observation consistent with the likelihood that Rv0077c is not a mono-oxygenase. It is possible that the effects of SMARt-402 is growth-condition dependent, that this molecule affects another pathway to either increase the susceptibility of *M. tuberculosis* to ETH, or that another SMARt-420-induced enzyme in addition to Rv0077c synergize together to activate ETH (A. Baulard, personal communication). Irrespective of these possibilities, it is unlikely that Rv0077c has a considerable role in the activation of ETH. While a previous report named Rv0077c and Rv0078, EthA2 and EthR2, respectively, we propose to rename them “LoaA” and “LoaR” for “<u>l</u>oss <u>o</u>f <u>a</u>cid-fast staining <u>A</u> and <u>R</u>epressor” due to the lack of association of these proteins with ETH susceptibility.

Finally, our studies have opened the door to the possibility that numerous commensal and pathogenic microbes (including fungi) use cytokinins for intra- or inter-species communication in complex systems such as the gut microbiome. The identification of one or more CK receptors and the signal transduction pathway that leads to the induction of Rv0077c expression will likely lay the foundation for understanding CK signaling in hundreds of bacterial species.

## Author Contributions

M.I.S. and K.H.D. performed *in vitro* and *in vivo M. tuberculosis* work. H.C.H. and H.L. determined the structure of Rv0078 and performed the mutagenesis and EMSA assays. M.B.J. performed the RNA-Seq analysis. S.H.B. and A.T.J. performed antibiotic susceptibility assays. M.S. provided cytokinins and their ribosides. V.J., M.R.M. and M.J. performed the lipid analysis. C.X. performed mass spectrometry of fatty acids extracted from purified Rv0078 protein. M.S. synthesized cytokinins and their precursors. M.I.S., H.C.H., H.L. and K.H.D. wrote the manuscript.

## Acknowledgments

We thank A. Darwin and V. Torres for critical review of a draft version of this manuscript. We thank S. Zhang for assistant in preparing reagents. This work was supported by NIH grant R01 HL092774 awarded to K.H.D. K.H.D. holds an Investigators in the Pathogenesis of Infectious Disease Award from the Burroughs Wellcome Fund. M.I.S. and S.H.B were supported by the Jan T. Vilcek Endowed Fellowship fund. S.H.B. is also supported by NIH grant T32 AT007180. H.C.H. and H.L. were supported by R01 AI070285. M. Strnad was supported by LO1204 from the National Program of Sustainability I. We thank A. Liang and Y. Deng of the Microscopy Laboratory at New York University Langone Medical Center for assistance with microscopy and imaging. Diffraction data for this study were collected at the Lilly Research Laboratories Collaborative Access Team (LRL-CAT) beamline and the Life Sciences Collaborative Access Team (LS-CAT) beamline at the Advanced Photon Source (APS), Argonne National Laboratory. APS was supported by the U.S. Department of Energy, Office of Science, Office of Basic Energy Sciences, under Contract DE-AC02-06CH11357. Use of the LRL-CAT beamline at Sector 31 of the APS was provided by Eli Lilly Co., which operates the facility.

## Materials and Methods

### Bacterial strains, plasmids, primers, chemicals, and culture conditions

Bacterial strains, plasmids, and primer sequences used in this study are listed in Table supplement 1. All primers for cloning and sequencing were from Invitrogen, Inc. *M. tuberculosis* strains were grown in Middlebrook 7H9 broth (Difco) supplemented with 0.2% glycerol, 0.05% Tween-80, 0.5% fraction V bovine serum albumin, 0.2% dextrose and 0.085% sodium chloride (“7H9c”). *M. tuberculosis* cultures were grown without shaking in 25 or 75 cm^2^ vented flasks (Corning) at 37 ºC. 7H11 agar (Difco) supplemented with 0.5% glycerol and BBL TM Middlebrook OADC enrichment (BD) was used for growth on solid medium (“7H11”). *M. tuberculosis* was transformed as described(Hatfull & Jacobs, 2000). *E. coli* strains used for cloning and expression were grown in LB-Miller broth (Difco) at 37 °C with aeration on a shaker or on LB agar. *E. coli* strains were chemically transformed as previously described(Sambrook *et al.*, 1989). The final concentrations of antibiotics used for *M. tuberculosis* growth: kanamycin, 50 µg/ml; hygromycin, 50 µg/ml; streptomycin, 25 µg/ml; and for *E. coli*: hygromycin, 150 µg/ml; kanamycin, 100 µg/ml; and streptomycin 50 µg/ml. AMP, adenine and iP were purchased from Sigma. iPR, 2MeSiP and 2MeSiPR were synthesized as previously described(Sugiyama & Hashizume, 1978). The purity of the synthesized cytokinin and derivatives was >98% for each as determined by high performance liquid chromatography and mass spectrometry.

The Rv0077c∷MycoMarT7 mutant was isolated from a library of ordered transposon insertion mutants as previously described(Darwin *et al.*, 2003, Festa *et al.*, 2007). The ΔRv0078∷*hyg* mutant was made by deletion-disruption mutagenesis as described in detail elsewhere using pYUB854(Festa *et al.*, 2011, Bardarov *et al.*, 2002).

### Protein purification and immunoblotting

DNA sequence encompassing the full-length Rv0078 or Rv0077c gene was cloned into pET24b(+) vector using primers listed in Table supplement 1. Recombinant proteins were produced in *E. coli* ER2566 and purified under native conditions for Rv0078 and denaturing conditions for Rv0077c according to the manufacturer′s specifications (Qiagen). Polyclonal rabbit antibodies were raised by Covance (Denver, PA). For all immunoblots, cell lysates or purified proteins were separated sodium dodecyl sulfate polyacrylamide gel electrophoresis (SDS-PAGE); transferred to nitrocellulose and incubated with rabbit polyclonal antibodies to the protein of interest at 1:1000 dilution in 3% bovine serum albumin in TBST (25 mM Tris-HCl, pH 7.4, 125 mM NaCl, 0.05% Tween 20, pH 7.4). Equal loading was determined by stripping the nitrocellulose membranes with 0.2 N NaOH for 5 min, rinsing, blocking and incubating the nitrocellulose with polyclonal rabbit antibodies to dihydrolipoamide acyltransferase (DlaT)(Tian *et al.*, 2005). Horseradish peroxidase conjugated anti-rabbit antibody (GE-Amersham Biosciences) was used for chemiluminescent detection (SuperSignal West Pico; ThermoScientific).

For crystallography studies of Rv0078-His_6_, bacteria were grown at 37°C to an OD_600_ = 0.5-0.6 before being induced with 0.5 mM IPTG and incubated at 16°C overnight. After harvesting by centrifugation, cells were lysed by passing through a microfluidizer cell disruptor in 10 mM potassium phosphate, pH 8.0, 10 mM imidazole, and 500 mM NaCl. The homogenate was clarified by spinning at 27,000 *g* and the supernatant was applied to a HiTrap-Ni column (GE Healthcare) pre-equilibrated with the lysis buffer. Histidine-tagged protein was eluted with a 10–300 mM imidazole gradient in 10 mM potassium phosphate, pH 8.0, containing 300 mM M NaCl. The Rv0078 fractions were further purified by a Superdex 75 column (16 × 1000 mm, GE Healthcare) pre-equilibrated with 20 mM potassium phosphate, pH 8.0, and 300 mM NaCl. The purified Rv0078 was concentrated to 40 mg/ml for crystallization screen.

### RNA-Seq and 5′RACE

Three biological replicate cultures of WT *M. tuberculosis* were grown to an OD_580_ ~1 and incubated in 100 µM iP or DMSO (control) for five hours. Cells were harvested and RNA was purified as described previously (Festa *et al.*, 2011). Briefly, an equal volume of 4 M guanidinium isothiocyanate, 0.5% sodium N-lauryl sarcosine, 25 mM trisodium citrate solution was added to cultures to arrest transcription. RNA was isolated with Trizol Reagent (Invitrogen) and further purified using RNeasy Miniprep kits and DNase I (Qiagen). Transcriptome profiling by RNA-seq was performed and analyzed as follows: RNA from *M. tuberculosis* cultures were extracted for library construction. Libraries were constructed and barcoded with the Epicentre ScriptSeq Complete Gold low input (Illumina, Inc) and sequenced on Illumina HiSeq 2000 sequencer using version 3 reagents. Unique sequence reads were mapped to the corresponding reference genome and RPKM values were calculated in CLC (CLC version 7.0.4). Genes with significantly different RPKM values were identified using the Significant Analysis for Microarray (SAM) statistical analysis component of MeV(Saeed *et al.*, 2003).

5′RACE was performed as described by the manufacturer (Invitrogen). Briefly, 1 µg of RNA was used as template for cDNA production using a reverse primer 150–300 bp downstream of annotated translational start sites. A 3′ poly-C tail was added to cDNA by recombinant Tdt. The cDNA was then amplified using a nested reverse primer and a primer that anneals to the poly-C tail. Products were cloned and sequenced. Likely transcriptional start sites were selected based on clones that had the most nucleotide sequence upstream of the start codon.

### EMSA

A series of double stranded DNA probes consisting of sequences in the intergenic region between Rv0077c and Rv0078 were generated by annealing two complementary oligonucleotides and a 5′-end IRDye700-labeled 14-nucleotide oligomers (5′dye-GTGCCCTGGTCTGG-3′) (Integrated DNA Technologies). Binding assays were performed by incubating 100 nM of probes and various concentrations of Rv0078 at room temperature for 30 min in 20 mM HEPES, pH 7.5, 3 mM DTT, 0.1 mM EDTA, 100 mM KCl, 5% glycerol, 5 mg/ml BSA, 10 mM MgCl_2_, and 0.25% Tween 20, and were subsequently resolved in 6% polyacrylamide gels in 0.5 × TBE buffer. Mobility shifts of protein-DNA complex were visualized in LI-COR Odyssey imager.

### Crystallization and structure determination

DNA-free Rv0078 crystals were obtained by screening at 20 °C using the sitting-drop vapor diffusion method. The C2 space group crystals were grown in 0.1 M sodium cacodylate, pH 6.4, and 1.3 M Lithium sulfate. SeMet substituted Rv0078 crystals with C2 space group were grown in in 0.1 M sodium cacodylate, pH 6.6, 1.3 M Lithium sulfate, 0.2 M magnesium sulfate, and 2% PEG400. Diffraction data to a resolution of 1.85 Å were collected at the Lilly Research Laboratories Collaborative Access Team (LRL-CAT) beamline of Advanced Photon Source (APS), Argonne National Laboratory, and were processed with Mosflm software(Winn *et al.*, 2011). The program Hybrid-Substructure-Search in the Phenix package was used to locate the Se sites and the initial phasing was carried out using the program Autosol of Phenix. The 2.7 Å map phased by SAD method allowed us to build Rv0078 model unambiguously. The native Rv0078 structure was subsequently determined by the program PHASER using SeMet substituted Rv0078 as the initial search model. To obtain the Rv0078-DNA crystals, purified Rv0078 was co-crystallized with a 23-mer DNA duplex (5′-TTTACAAGCAGACTGCCGGTAAC-3′) at a molar ratio of 2:1 (protein dimer:DNA) in the presence of 150 mM MgCl_2_. The DNA-bound Rv0078 crystals were grown in the buffer containing only 0.2 M Magnesium formate. Diffraction data to 3.0 Å were collected at the Life Sciences Collaborative Access Team (LS-CAT) beamline of APS and were processed with Mosflm. The Rv0078-DNA structure was determined by PHASER using DNA-free structure as the search model. All the refinements were performed using Phenix-refine (Adams *et al.*, 2010). The statistics were provided in **Supplemental Table 3**.

### Mass spectrometry of fatty acids co-purified with Rv0078

Fatty acids were extracted from 5 mg Rv0078 that was purified from *E. coli*, and analyzed as described previously (Fan *et al.*, 2013). Separation and identification of the FA methyl esters were performed on an HP5975 gas chromatograph-mass spectrometer (Hewlett-Packard) fitted with a 60 m × 250 µm SP-2340 capillary column (Supelco) with helium as the carrier gas

### Isothermal calorimetry

ITC experiment was performed in a Microcal PEAQ-ITC at 25 °C. The stirring speed was 750 rpm and the interval between each titration was 150 sec. The concentration of Rv0078 in the reaction cell was 25 µM and the concentration of the titration DNA ligand was 500 µM. The recorded thermal data was analyzed using Microcal PEAQ-ITC analysis software.

### MIC_50_ Determination

To determine the MIC_50_ of each antibiotic, *M. tuberculosis* strains were grown to an OD_580_ ~0.7 and diluted into fresh media to an OD_580_ of 0.02. Diluted cultures were transferred to a 96-well microtiter plate containing triplicate, 10-fold serial dilutions of antibiotic. Cell wall-active antibiotics (vancomycin, meropenem) were supplemented at all concentrations with potassium clavalunate to inhibit the intrinsic β-lactamase activity of *M. tuberculosis*. After five days of incubation at 37ºC, growth of each strain was measured by OD_580_. MIC_50_ values were interpolated from a non-linear least squares fit of log_2_-transformed OD_580_ measurements. Data are representative of two independent experiments. Antibiotics were purchased from Sigma-Aldrich (clavulanate, ethionamide, meropenem, rifampicin, vancomycin) or Thermo-Fisher Scientific (ciprofloxacin, ethambutol, isoniazid, norfloxacin, streptomycin).

### Mouse Infections

Mouse infections were performed essentially as described previously(Darwin *et al.*, 2003). 7-9-week-old female C57BL6/J mice (The Jackson Laboratory) were infected by aerosol to deliver ~200 bacilli per mouse, using a Glas-Col Inhalation Exposure System (Terre Haute, IN). Strains used were WT (MHD761) or (MHD794), Rv0077c (MHD1086), Rv0077c complemented (MHD1077), Rv0078 (MHD1315), Rv0078 complemented with WT Rv0078 (MHD1318) and Rv0078_W100R_ (MHD1316). This study was performed in strict accordance with the recommendations in the Guide for the Care and Use of Laboratory Animals of the National Institutes for Health. Mice were humanely euthanized according to an approved Institutional Animal Care and Use Committee protocol at New York University School of Medicine. Lungs and spleens were harvested and homogenized PBS/0.05% Tween-80 at indicated time points to determine bacterial CFU.

### Metabolomic analysis of *M. tuberculosis* cell lysates

Four independent cultures of each analyzed strain were grown in 7H9 to an OD_580_ ~0.7 and treated with iP in DMSO at a final concentration of 100 µM or an equal volume of DMSO for 24 hours. Bacteria were harvested the next day at OD_580_ ~1. Sixty-five OD-equivalents per replicate were processed by chloroform:methanol extraction(Layre *et al.*, 2011, Samanovic *et al.*, 2015). Metabolomic profiling was performed by Metabolon, Inc.

### Staining and microscopy

*M. tuberculosis* strains were grown to mid-logarithmic phase (OD_580_ ~0.5-0.7). 5 µl of culture was spotted onto glass slides and heat-killed over a flame or on a heat block (15 min, 80°C). Staining was performed according the method of Ziehl-Neelson as per the manufacturer’s instructions (BD Stain Kit ZN). Images were acquired on a Zeiss Axio Observer with a Plan-Aprochromat 63x/1.4 oil lens. Images were taken with an Axiocam503 camera at the NYULMC Microscopy Laboratory.

### Analysis of total lipids, mycolic acids and shorter chain fatty acids

For lipid analysis 400 ml cultures were grown up to OD_580_ ~0.7 and treated with iP in DMSO at a final concentration of 100 µM or an equal volume of DMSO-only for 24 hours. 400 ml of Rv0077c and Rv0078 cultures were treated with DMSO only. Cells were washed three times in DPBS and heated at 100°C for 45 min for sterilization before freezing at −20 °C.

Total lipids extraction from bacterial cells and preparation of fatty acid and mycolic acid methyl esters from extractable lipids and delipidated cells followed earlier procedures (Stadthagen *et al.*, 2005). Total lipids and fatty acid/mycolic acid methyl esters were analyzed by one and two-dimensional thin-layer chromatography (TLC) in a variety of solvent systems on aluminum-backed silica gel 60-precoated plates F_254_ (E. Merck). TLC plates were revealed by spraying with cupric sulfate (10% in a 8% phosphoric acid solution) and heating. Alternatively, total lipids were run in both positive and negative mode and the released fatty acids/mycolic acids in negative mode only, on a high resolution Agilent 6220 TOF mass spectrometer interfaced to a LC as described(Sartain *et al.*, 2011, Bhamidi *et al.*, 2012). Data files were analyzed with Agilent’s Mass hunter work station software and most compounds were identified using a database of *M. tuberculosis* lipids developed in-house (Sartain *et al.*, 2011). Fatty acids methyl esters from extractable lipids were treated with 3 M HCl in CH_3_OH (Supelco) overnight at 80°C, dried and dissolved in *n*-hexane(s) prior to GC/MS analysis. GC/MS analyses of fatty acid methyl esters were carried out using a TRACE 1310 gas chromatograph (Thermo Fisher) equipped with a TSQ 8000 Evo Triple Quadrupole in the electron impact mode and scanning from *m/z* 70 to *m/z* 1000 over 0.8 s. Helium was used as the carrier gas with a flow rate of 1 ml per min. The samples were run on a ZB-5HT column (15 m × 0.25 mm i.d.) (Zebron). The injector (splitless mode) was set for 300°C (350°C for mycolic acid methyl esters). The oven temperature was held at 60°C for 2 min, programmed at 20°C per min to 375°C, followed by a 10 min hold. The data analyses were carried out on Chromeleon data station.

